# Experimental assessment of marine microbial interactions: from predatory protists promoting bacterial survival to bacterial lysis of the protists

**DOI:** 10.1101/2024.02.09.579682

**Authors:** Diana Axelsson-Olsson, Nikolaj Gubonin, Stina Israelsson, Jarone Pinhassi

## Abstract

Bacteria in aquatic environments are a principal food source for predatory protists. Whereas interactions between bacteria and protists are recognized to play important roles in determining the pathogenesis and epidemiology of several human pathogens, few studies have systematically characterized the interactions between specific aquatic bacteria and protists beyond the prey-predator relation. We therefore surveyed the outcome of individual co-cultures between 18 different genome-sequenced marine bacteria with known virulence gene repertoires and three model protist species widely used for assessing bacteria-protist interactions. Strikingly, ten, five, and three bacterial isolates were capable of lysing the protists *Acanthamoeba polyphaga, Tetrahymena pyriformis* and *Euglena gracilis*, respectively. A majority of the bacteria were able to grow and/or maintain viable populations in the presence of viable protists. Some bacteria survived longer in the presence of viable protists but not heat-killed protists, and were observed in protist vacuoles. In this respect, thus, marine bacteria are similar to several protist-dependent human pathogens, including *Legionella*. Analyses of growth patterns in low-nutrient media showed that co-cultivation with *A polyphaga* allowed one bacterial strain to overcome nutritional stress and obtain active growth. Five isolates depended on viable amoebae to grow, notwithstanding nutrient media status. The remarkable capability of surviving encounters with, and even actively killing, bacterivorous protists, indicates that diverse (and possibly novel) bacterial defense strategies and virulence mechanisms to access nutrients are widespread among marine bacteria. The diversity of interactions uncovered here has important implications for understanding ecological and evolutionary consequences of population dynamics in bacteria and protists.

**IMPORTANCE:** The microbiome constitutes the base of food webs in marine waters. Its composition partly reflects biotic interactions, where bacteria primarily are considered as prey of predatory protists. However, studies that focus on one or a few species have shown that some bacteria have abilities to escape grazing and may even be capable of lysing their protist predators. In this study, we substantially extend these findings by systematically investigating interactions among multiple taxa of both bacteria and protists. Our results show that marine bacteria display a wider and more complex range of interactions with their predators than generally recognized - from growth dependency to protist lysis. Given that such interactions play key roles in the pathogenesis and epidemiology of several human pathogens, our findings imply that bacterial virulence traits can contribute to defining the structure and ecology of the marine microbiome.

## INTRODUCTION

Bacteria constitute the basis for microbial food webs in the aquatic environment, being the principal food source for predatory unicellular eukaryotic organisms - protists - like heterotrophic flagellates. In most environments, however, in addition to the bacteria being grazed by bacterivores, bacteria and protists interact in multiple ways as a consequence of a very long shared evolutionary history. Interkingdom interactions between these microbes include parasitism, mutualism, and symbiosis (1). From the bacterial perspective this can result in prolonged survival, enhanced dissemination, gene transfer, and even altered pathogenicity (2–5). Some bacteria are resistant to protist grazing (6, 7) and may in fact produce toxic compounds that are lethal for heterotrophic and phototrophic protists (8–12). Hence, interactions between these microorganisms could be an important factor of the ecology of the ocean, not only from a grazing perspective, but also from a perspective where bacteria could control protist viability and abundance. For example, it has been proposed that algicidal bacteria can be involved in terminating algal blooms (13). The mechanisms behind such potential effects are largely unknown and knowledge on prevalence and function of interactions between bacteria and protists in marine environments, other than the latter preying on the former, remain limited.

From a human health perspective, in contrast, the mechanisms behind different types of interactions between bacteria and protists are well studied, since they in several cases are crucial for bacterial pathology and epidemiology (reviewed by Greub and Raoult (2004) (14) and Balczun et al. (2017) (15)). As a prime example, co-culture experiments of the human pathogen *Legionella pneumophila* with the amoeba *Acanthamoeba castellanii* show that bacterial survival and dissemination critically depend on the association of bacteria with the amoeba. Moreover, the interaction also enhances the ability of the bacteria to invade macrophages (3) and augments intracellular survival in monocytes (16). Also, for the pathogen *Mycobacterium avium* the virulence against human macrophages is significantly enhanced by replication in *A. castellanii* (17). In a similar manner, interactions with the amoeba *A. polyphaga* elevate cytotoxicity and pro-inflammatory potential of bacteria in murine macrophages (18). However, the opposite situation is observed for *Francisella novicida* where the interaction with *A. castellanii* results in a decreased infectivity of mice (19). These studies show that, in addition to bacteria being grazed by bacterivores, microbial interactions between protists and bacteria may determine the pathogenicity and epidemiology of bacteria.

Similar to the medically important amoebae examples above, unicellular eukaryotes in aquatic environments may be potential promoters of bacterial survival, persistence and growth, possibly offering shelter, site for reproduction and increased dissemination (2, 20–23). Currently, novel culturing strategies and high-throughput sequencing analyses are casting light on the extensive phylogenetic diversity and phenotypic characteristics of protists dominant in the sea and in freshwater. Pathogenic bacteria, capable of benefiting from interactions with protists (24, 25), have been found in fish and aquatic environments (26). Such strains may contribute to serious economic losses in marine and freshwater ecosystems, impacting both fish industry and marine recreation activity, as well as possibly being a threat to human health.

Earlier studies have indicated that microbial interactions between bacteria and protists in aquatic environments can be complex and that specific interactions depend on the species pairs under study (8). To exemplify, marine diatom interactions with bacteria are directly linked to the species composition of both algal and bacterial assemblages and to the physiological state of the algae (27, 28). However, the prevalence, long-term effects, and molecular mechanisms of interactions between free-living bacteria and protists are not well known. One contributing factor to a multitude of interaction modes could be bacterial expression of virulence genes. As shown by Persson et al. (2009) (29), virulence gene homologues are abundant in marine bacteria. The functions of such genes in marine bacteria are generally unknown, but their presence indicates that some bacteria may use them to infect or consume eukaryotic cells such as protists. Indeed, this has been shown to be the case with *Pseudomonas aeruginosa*, which uses type III secretion system to kill biofilm-associated amoebae (30). The virulence of this system, with a particular focus on aquatic pathogens, is reviewed by Rahmatelahi et al. (2021) (31). To better understand the implications of interactions between bacteria and protists in the ocean, knowledge on the prevalence and nature of these interkingdom relationships is necessary.

The aim of the present study was to survey the interactions between bacteria and protists during long-term co-culturing. Often studies investigate interactions of one or a few model bacteria or protists using laboratory-specific experimental setups, which makes comparisons between studies or taxa difficult. We did a survey to systematically investigate 18 different marine bacterial species with defined virulence gene repertoires (29) and three widely studied model protist species, which are found in various aquatic environments, to evaluate the variety of possible interactions. The protists included in the study were *A. polyphaga*, a phagocytic amoeba, well known to harbor bacteria which benefit from this interaction (14, 21, 32); *Euglena gracilis,* a mixotrophic flagellate; and *Tetrahymena pyriformis,* a ciliate with high grazing capability (33). All three protists were previously examined in an interaction study with the food borne pathogen *Campylobacter jejuni* (34). Due to the prominent position of amoeba as widely used model protists for interactions with bacteria, interactions with *A. polyphaga* were investigated in furthest detail. Our null hypothesis was that co-culturing of bacterivorous protists with bacteria would result in equal inhibition of growth of the studied bacteria due to grazing. On the contrary, our results showed that several bacterial species were potent killers of protists, and in several cases viable protists increased bacterial survival capability.

## RESULTS

### Selective and density dependent bacterial-mediated lysis of protists

Several bacterial species were observed to have negative effects on survival of the protists. The ciliate *T. pyriformis* was lysed as an effect of co-culturing with five different bacteria at an initial inoculation ratio of bacteria:protists of 10:1 (Table 1). The flagellate *E. gracilis* was lysed by three different bacterial species (at both high and low inoculation ratio). Interestingly, co-culturing with the bacterial strains BAL199 and AND4 caused decreased and increased green colouration on *E. gracilis*, respectively. Seven bacterial strains caused *E. gracilis* to alter its morphology to a more rounded shape. Ten different bacteria lysed the amoeba *A. polyphaga* when added in the high bacteria:protist ratio (Table 1). Four of these bacteria were also able to lyse the amoeba cells when the bacteria were added in a much lower ratio (1:1000). Additionally, four bacteria altered the amoebae morphology when added in the high bacteria:protist ratio (10:1), from typically stretched out trophozites to rounded cells typical for stressed amoeba.

**Table 1.**
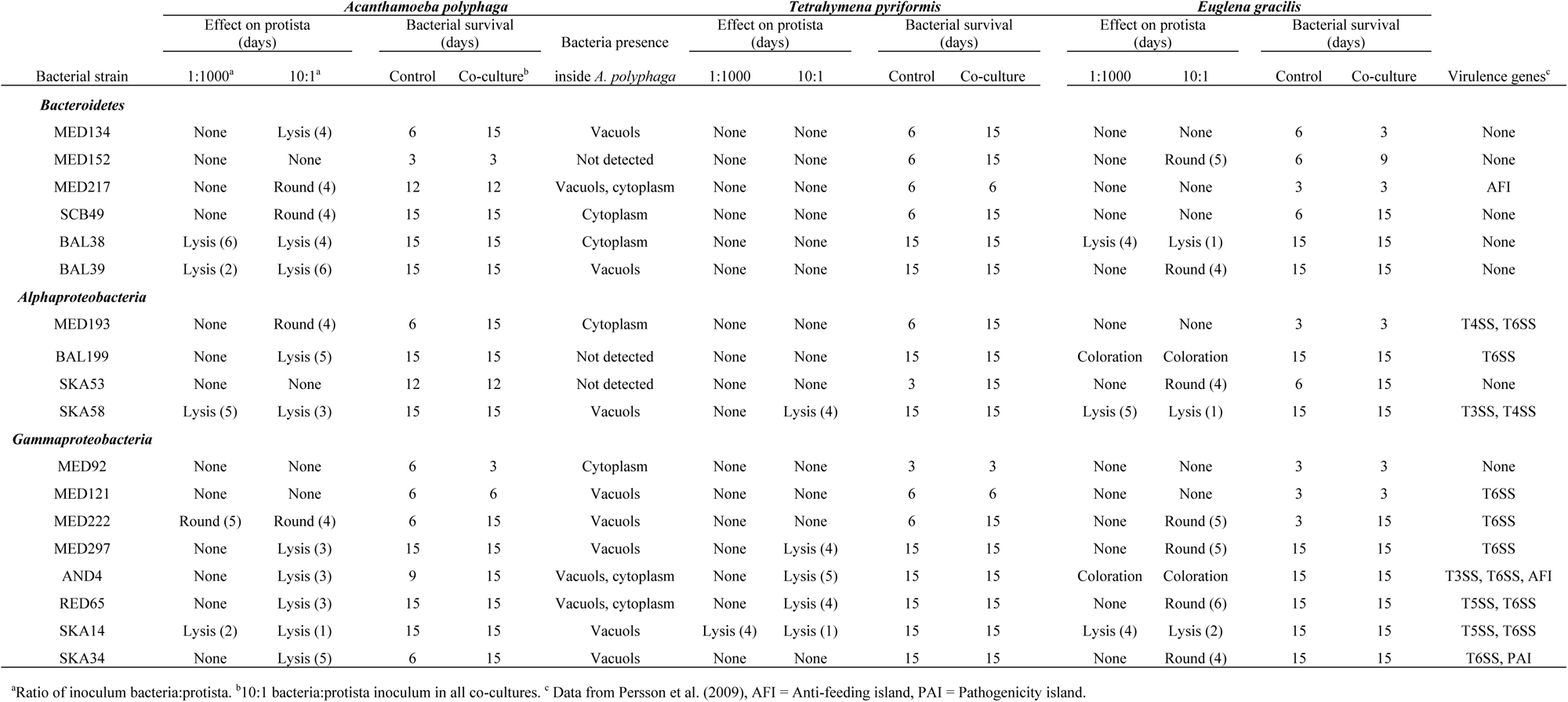
Marine bacterial species included in the study and their interactions in co-cultures with *Acanthamoeba polyphaga*, *Tetrahymena pyriformis* and *Euglena gracilis*. Survival in co-culture, bacteria-induced lysis and other effects, intra-amoebal presence of bacteria in *A. polyphaga* and the virulence gene repertoire for the bacterial strains are shown.

There were notable differences between bacteria in the ability to lyse the model protists. The bacterium *Stenotrophomonas* sp. SKA14 (Gammaproteobacteria) was the only species that lysed all three protists (within one to four days), regardless of the inoculum ratio (Table 1). As seen in Fig. 1, SKA14 caused *A. polyphaga* to start forming cysts and lyse within 24 hours and eventually causing complete lysis within one week, while growing to high bacterial concentrations. Also, *Sphingomonas* sp. SKA58 (Alphaproteobacteria) was capable of lysing all three protists, although less potently. Co-culturing of *A. polyphaga* and *T. pyriformis* with several Gammaproteobacteria (i.e., MED297, AND4 and RED65) at high inoculum ratio (10:1) caused these protists to lyse, while these bacteria only caused changes in cell morphology of *E. gracilis* (Table 1). BAL38 (Bacteroidetes) on the other hand, caused *A. polyphaga* and *E. gracilis* to lyse during co-culture starting with both a high and a low inoculum, while *T. pyriformis* did not indicate any co-culturing effect with this bacterial species.

**Fig 1.**
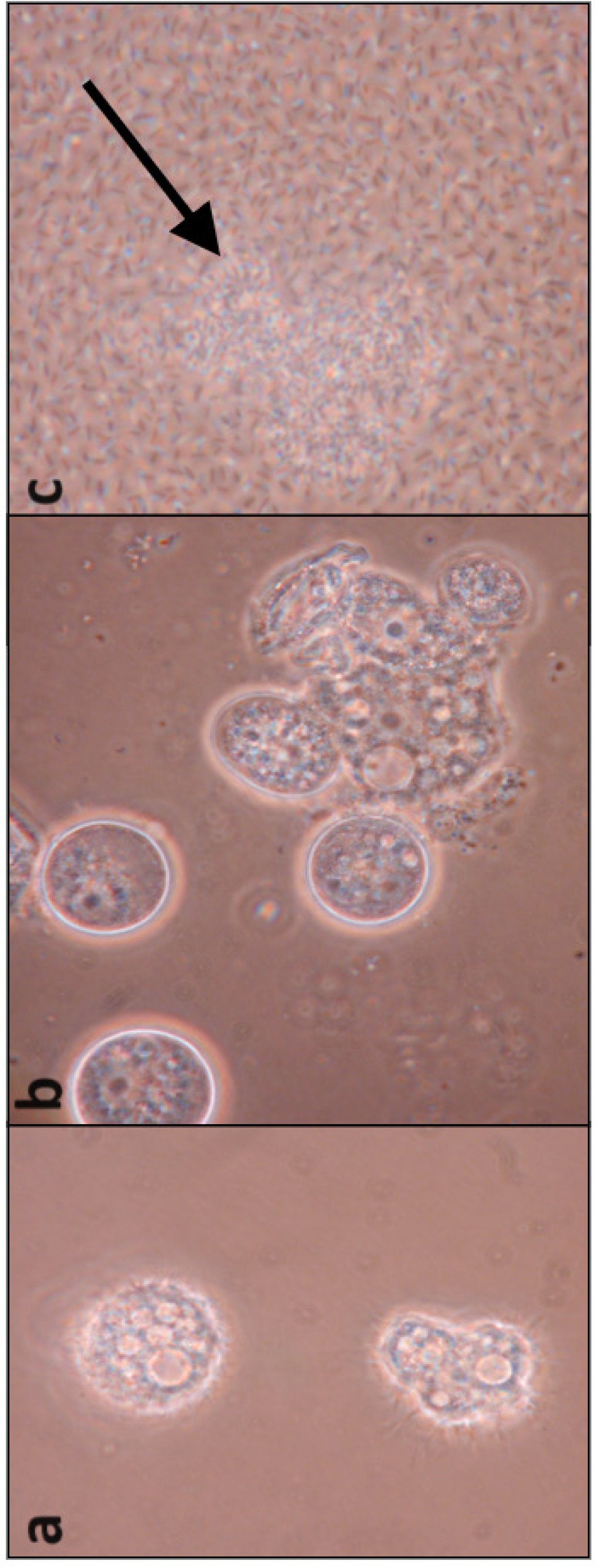
Microscope images of *Stenotrophomonas* sp. strain SKA14 and amoebae in co-culture. (**a**) 1 hour incubation, low bacterial concentration, amoebae in trophozoite form. (**b**) 24 hours incubation, moderate bacterial concentration, amoebae start forming rounded cysts. (**c**) One week incubation, rich bacterial concentration, ruptured amoebas (arrow).

### Enhanced or maintained bacterial survival in co-culture with protists

Bacterial survival was in several cases stimulated by co-culturing (Table 1). Also, many bacteria did not show any differences in survival time when comparing protist co-culturing with bacterial monoculturing. Only in two cases did the protists (*A. polyphaga and E. gracilis*) degrade all the bacteria in co-culture (MED92, MED134). Most of the bacteria survived through the entire 15-day experiments. Co-culturing with *T. pyriformis* resulted in a prolonged survival for six bacterial strains (MED134, MED152, SCB49, MED193, SKA53 and MED222) as compared to bacterial survival in medium alone. *E. gracilis* caused a prolonged bacterial survival in four cases (MED152, SCB49, SKA53 and MED222).

Bacterial co-culturing with *A. polyphaga* prolonged the survival of five bacterial species (MED134, MED193, MED222, AND4 and SKA34). MED222 was the only bacterium that showed a prolonged survival when co-cultured with all three protist species, as compared to culturing in protist medium alone. During this experiment bacterial presence within *A. polyphaga* cells was also assessed. The results showed that 15 out of 18 bacterial species (all but MED152, BAL199 and SKA53) were detected inside amoeba cells, either within vacuoles or in the cytoplasm, or in some cases both (Table 1).

Having found that bacteria and protist co-culturing may result in very different effects (Table 1), interactions between bacteria and *A. polyphaga* were investigated in further detail (Fig. 2, Table S1). The importance of amoebae viability was investigated, by comparing co-cultures using both viable and heat-inactivated amoebae. Additionally, possible interactions from heat-inactivated previously viable co-cultures, was also investigated. Fig. 2 shows the different outcomes, represented by the dynamics of seven of the bacterial species, observed with all 18 bacterial strains. Strikingly, SKA34, MED222, AND4, MED134 and MED193 all benefited the most from the presence of viable but not dead amoebae, in comparison to bacterial monocultures. The *Photobacterium* sp. SKA34 for example, had a maximum survival time of 19 days in the presence of viable amoeba cells, compared to only one to six days when cultured without viable amoebae. Other bacteria did not survive any treatment, as represented by MED152, or tolerated all conditions well as represented by SKA14 (Fig. 2). Survival times of all strains is shown in Table S1.

**Fig 2.**
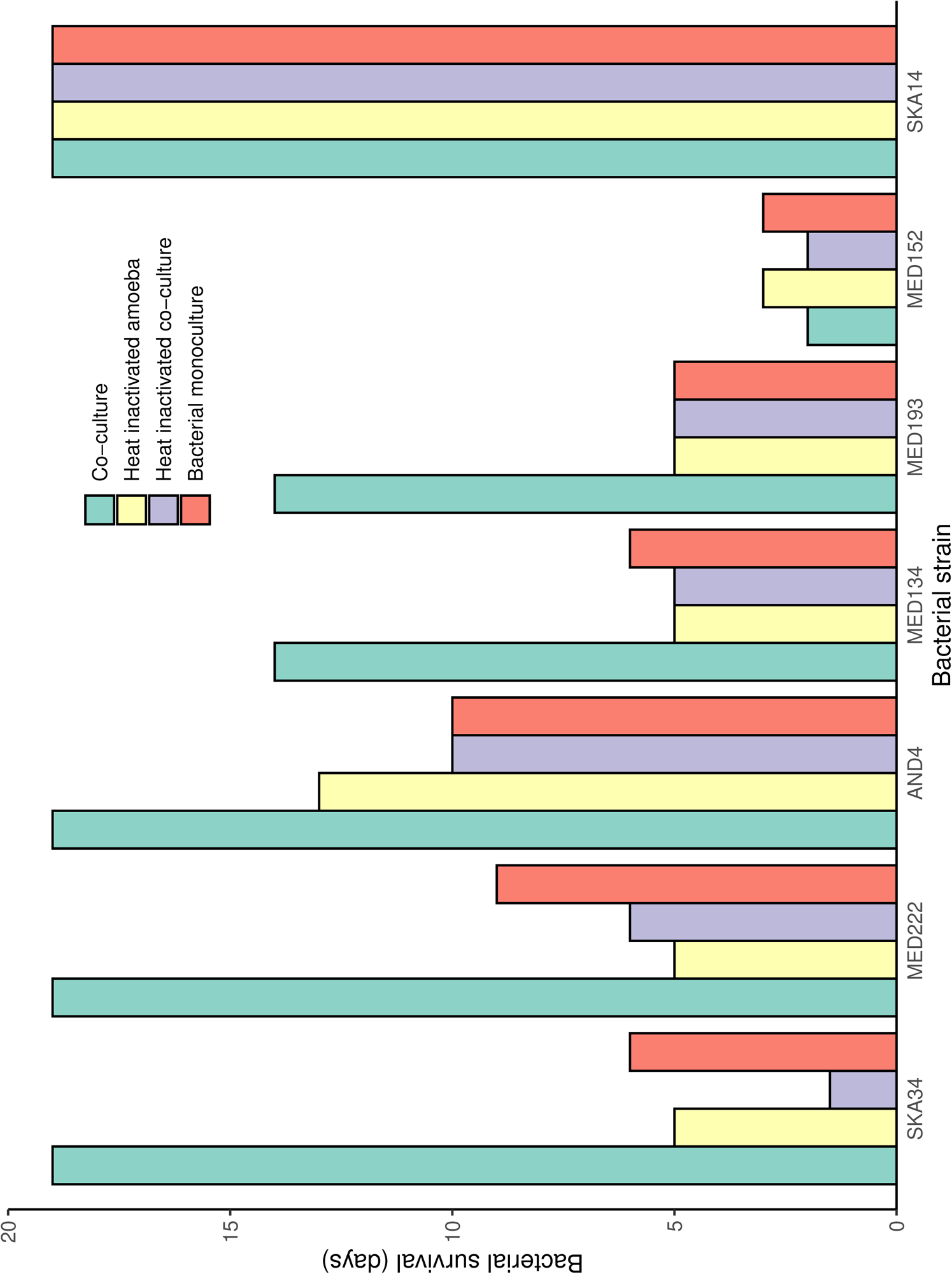
Bacterial survival in co-culture depending on amoeba viability. Bacterial strains were cultured in the presence of viable *A. polyphaga* (green bars), heat inactivated *A. polyphaga* (yellow bars) or a heat inactivated previous co-culture of *A. polyphaga* and the same bacterial strain (purple bars) and compared to bacteria growing in monoculture (red bars). Strains that showed a dependence on amoeba viability (SKA34, MED222, AND4, MED134 and MED193) are shown as well as two examples of independence of amoeba viability in co-culture (MED152 and SKA14). Survival time of all strains included in this study are shown in Table S1. Duplicate samples were incubated for a maximum of 19 days. Bacterial viability was tested by plating, initially after 24 hours followed by every second day, and shown as mean values.

### Bacterial co-culture with *A. polyphaga* under different nutrient conditions reveal diverse dynamics

The effect of nutrient status of culture media on the amoeba-bacteria interactions was investigated by co-culturing in nutrient-rich medium (PYG) and low-nutrient medium (PBS) using bacterial inocula from late exponential growth phase. Monoculturing in PBS did not support neither bacterial nor amoebal growth (although *A. polyphaga* is known to survive for weeks in PBS), as shown for representative bacterial strains in Fig. 3.

**Fig 3.**
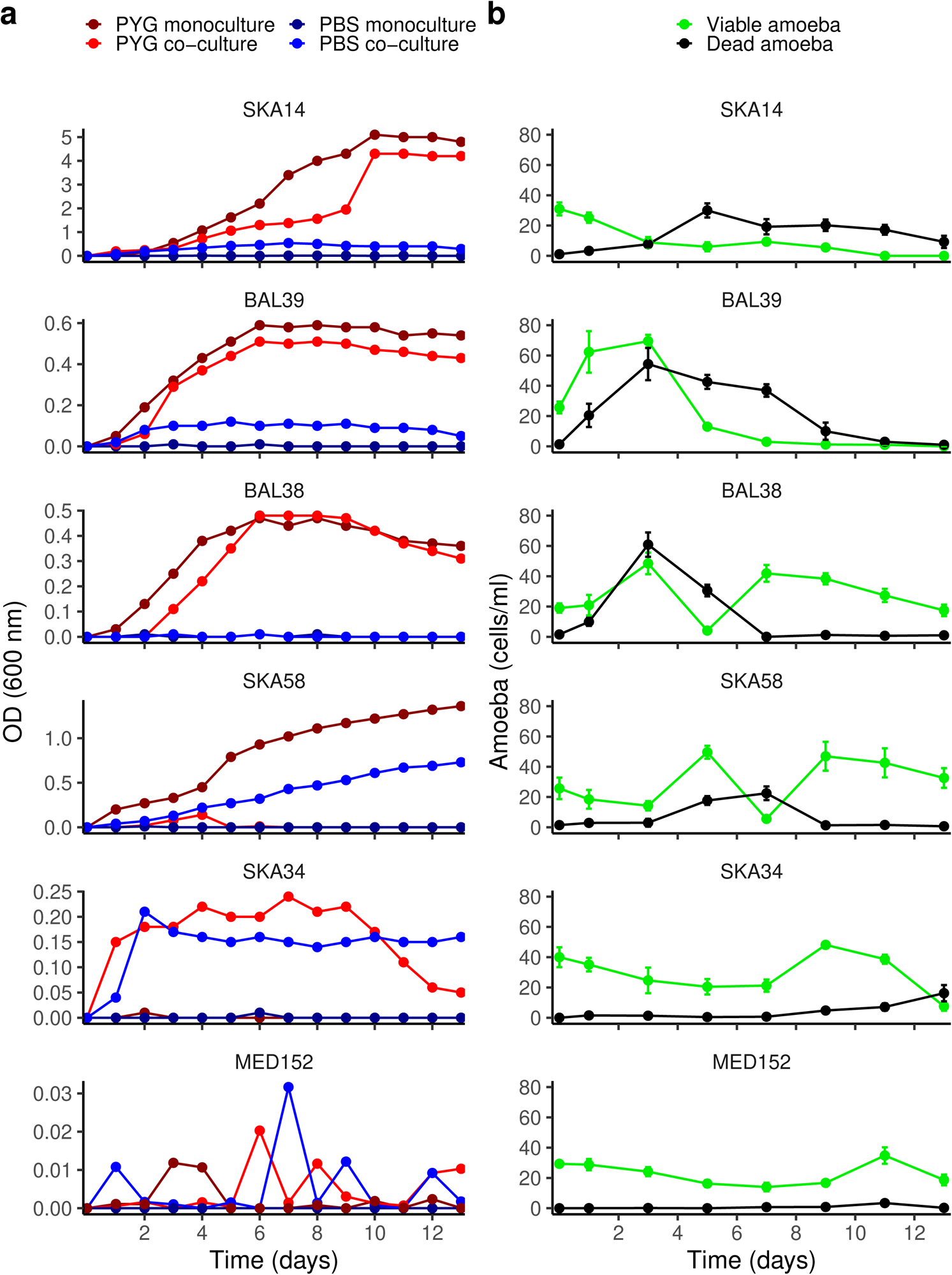
Growth curves of six bacterial strains and *A. polyphaga* viability in co-cultures with respective bacteria. (**a**) The bacteria were cultivated in nutrient-rich media (peptone yeast glucose [PYG]), low-nutrient media (phosphate buffered saline [PBS]), and in mono- and co-culture with *A. polyphaga*. (**b**) *A. polyphaga* viability in co-cultures with each of the bacterial strains. Note the different y-axis scales of the bacteria growth curves. All bacteria cultures started from late exponential growth phase. Bacterial growth and amoeba viability was determined by plating and trypan blue exclusion, respectively. For amoeba viability, error bars display the standard deviation across two replicates. Bacterial growth curves of the remaining 12 strains included in the study are shown in Fig. S1.

SKA14 reached high optical density values (approximately 4-5), when grown in PYG medium for 13 days (Fig. 3a). When grown in PYG co-culture with amoebae, growth was slower for the first 8 days, but from day 10 the OD reached around 4. As bacterial growth progressed, the viability of *A. polyphaga* decreased, and when the bacteria reached stationary phase on day 10, all amoebae had died (Fig. 3b). SKA14 did not grow in PBS alone but showed a minor increase in OD when co-cultured with amoebae in PBS (Fig. 3a).

BAL39 interacted with the amoeba in a similar way as SKA14, but differed in that it reached lower maximum OD-values, around 0.6, after 6 days, and in that growth was only slightly inhibited by co-culturing with *A. polyphaga* (Fig. 3a). Although BAL39 reached lower OD values, its growth led to death and subsequent disintegration of nearly all *A. polyphaga* cells by day 8 (Fig. 3b). In PBS, co-culturing with amoebae generated a modest growth of BAL39, while no growth was observed in PBS without amoebae (Fig. 3a).

BAL38 growth in co-culture with amoeba in PYG was delayed by approximately 1 day as compared to bacterial monoculture in PYG, although both cultures reached OD maxima at nearly 0.5 (Fig. 3a). Although amoeba fitness was clearly negatively affected by co-culturing with BAL38 initially, as shown by a simultaneous increase and then decrease in abundance of both viable and dead amoebae, the amoebae recovered as compared to the amoebae monoculture control (Fig. 3b and 3a, respectively). The BAL38 bacteria did not grow in PBS, either with or without amoeba.

SKA58 reached an OD value of approximately 1.4 when grown for 13 days in PYG (Fig. 3a). When grown in co-culture with amoeba in PYG media, slow growth to only OD=0.2 until day 4 was observed, after which the amoeba grazing cleared the cultures of bacteria. The amoeba fitness during the co-incubation in PYG was first positively affected, followed by a drastic viability decline, where the abundance of dead amoebae increased simultaneously, from which the amoebae subsequently managed to recover (Fig. 3b). While SKA58 did not grow in PBS alone, the bacteria grew well in PBS with amoeba, reaching an OD of about 0.7 by the end of the experiment.

SKA34 was not capable of growth in neither PYG nor PBS (Fig. 3a). However, in co-culture with *A. polyphaga* in both PYG and PBS, SKA34 rapidly grew to OD values around 0.2. Thereafter, ODs remained stable in PBS, but decreased towards the end of the experiment in PYG. The viability of the amoebae was not substantially affected by co-culturing with SKA34 (Fig. 3b), as compared to the amoebae monoculture control (MED 152, Fig. 3b). Similar dynamics in OD values in PBS and PYG were also observed with the bacterial strains MED222, AND4, MED217, MED92 and MED193 although MED92 and MED193 grew more poorly in co-culture in PBS compared to PYG (Fig. S1).

As exemplified by MED152, some bacterial species did not grow neither in PYG nor PBS, irrespectively of amoeba co-culturing (Fig. S1). Similar dynamics were also observed for MED121, MED134, MED 297, SCB49, SKA53, RED65 and BAL199, although in the case of MED134 some growth but poor survival was observed in amoeba co-culture in PYG (Fig. S1).

In summary, nine bacterial strains were capable of growth in both PYG and PBS when cultured together with viable (and presumably grazing) amoeba cells. Two of these bacterial strains (SKA14 and BAL39) were also capable of killing the amoeba cultures completely.

## DISCUSSION

The current work was inspired by studies in medical microbiology where analyses of interactions between bacteria and model unicellular eukaryotes - protists - have provided insights into how microbial interactions influence bacterial pathogenesis and epidemiology (15, 34–38). The potential ecological importance of interactions between bacteria and protists in a marine context, other than bacteria being a primary source of food for protists, is being increasingly acknowledged (8, 39, 40). We surveyed the outcome of interactions between a wide variety of phylogenetically distinct marine bacteria and three different protists under defined experimental conditions, allowing direct comparisons of their performance. This showed what types of interactions occur, how widespread and specialized they may be, and if any of the marine bacteria could gain fitness benefits from interacting with protists. All together, we observed four modes of interactions (beyond bacteria being prey for protists): *(i)* bacteria lysed the protists, *(ii)* simultaneous maintenance of bacterial and protist viability, *(iii)* bacterial utilization of protists as a source of nutrients under low-nutrient conditions, and *(iv)* viable protists enhancing, or even being necessary for, bacterial growth.

### Virulent bacteria display selective and density dependent lysis of protists

Lysis of protists during bacterial growth was observed with ten of the studied bacteria. Our survey revealed pronounced differences in how vulnerable the three protists were to bacterially mediated lysis, with *A. polyphaga* being most sensitive, and *T. pyriformis* and *E. gracilis* also differing in their modes of responses. For example, *E. gracilis* showed stress responses by rounding up or by altering its pigmentation. It is unclear why pigmentation increased in response to bacteria exposure in some cases. Previous studies have identified algicidal compounds produced by marine bacteria that on the contrary decrease pigmentation, followed by cell rupture, of various algae species. These reactions are possibly connected to increased levels of reactive oxygen species (10, 11). Because *E. gracilis* has a mixotrophic lifestyle, it may not be as sensitive to toxic extracellular compounds that targets the photosynthesis system (10), which could, in part, explain its atypical reaction.

In the cases where lysis of protists was observed, bacterial lysis efficacy was highest in the treatment with the higher bacteria:protist inoculation ratio. This efficacy was displayed both with respect to if lysis occurred or not and with respect to the time it took until lysis (with longer time required for co-cultures with low initial bacteria:protist ratio). The bacterium SKA14 (Gammaproteobacteria) showed the broadest lysis ability, being capable of killing all three protists included in the study at both high and low bacteria:protist inoculation ratios. A more limited lysis ability was displayed by for example BAL39, which only lysed *A. polyphaga*. Still, BAL39 was highly efficient in lysing the amoeba, as seen from its ability to lyse amoebae at both high and low bacteria:protist ratios. Our survey shows that lysis efficacy on particular protists varies considerably between bacteria in a species-specific manner, and that bacteria show distinct differences with respect to the range of protists they can lyse. Bacteria growing in high densities may form biofilms that are typically more tolerant to antimicrobial agents (41) and may result in cytotoxic effects on protist grazers (22). Other bacteria found within aquatic environments have also been shown to lyse protists (10, 20, 21, 42). Our results generally agree with the findings in these reports that lysis occurs within 3 - 4 days of co-cultivation with pathogenic bacteria. Similarly to strain-specific grazing of protists (43), predatory bacteria too may display strain-specific lysis of protists (9, 42).

The mechanism of bacterial-mediated lysis of protists differs, where some bacteria are reproducing intracellularly (20, 21, 44, 45) and disrupting intracellular components (46) until the protist bursts. In other taxa, lysis occurs through extracellular toxic compounds secreted by the bacteria (10). The latter seems to be the preferred mode with algicidal bacteria, although lysis as a result of direct contact of algicidal bacteria also occurs (13). Potential virulence mechanisms in our study are discussed in more detail below.

### Escaping grazing results in long-term co-viability of bacteria and protists

Simultaneous long-term protist viability and bacterial survival was observed with 14 of the 18 studied bacteria with at least one of three protists; 24 out of 54 interactions in total, of which four bacterial strains displayed intracellular presence within *A. polyphaga*. Indeed, aquatic prokaryotes have developed numerous strategies to avoid predation (47). For example, bacteria may take “shelter” within cellular compartments of the amoeba, protecting them from degradation (22). Alternatively, viable bacteria can be detected in co-cultures because they manage to survive at abundances below a threshold at which grazing is efficient. Indeed, despite efficient initial grazing of marine bacteria by *A. castellanii*, bacteria could still be identified 48 hours post infection, although at low levels (22). These findings on the ability of bacteria to survive for several days to weeks together with protists indicate that mechanisms for escaping protist grazing are widespread among marine bacteria.

#### An example of bacterium growth stage affecting amoebal viability in a complex way

A very complex example of coinciding protist and bacterial viability, which could be considered a mix between protist lysis by bacteria and simultaneous protist viability and bacterial survival was observed in co-cultures of BAL38 and *A. polyphaga*. This bacterium was capable of lysing both *A. polyphaga* and *E. gracilis*, but the growth curve experiments with amoeba co-cultures showed that the amoeba transiently recovered when BAL38 reached stationary growth phase. Thus, bacterial growth was initially delayed by amoeba grazing, but eventually bacterial growth would affect amoeba viability negatively. Finally, once the bacterial stationary phase was reached, the amoebae were given an opportunity to recover. This indicated that BAL38 can escape from predation by protists and may be virulent to such species during proliferation, even though they may coexist when the bacteria are in stationary phase. This in some respects mirrors the ‘Jekyll-and-Hyde’ relationship between the coccolithophore *Emiliania huxleyi* and its symbiotic bacterial partner *Phaeobacter gallaeciensis*. *P. gallaeciensis* first promotes the growth of *E. huxleyi*, only to transition into a pathogenic phase once *E. huxleyi* reaches higher densities (48). Similarly, a *Microbulbifer* strain showed increased algicidal effects against various algae species when at stationary phase as compared to exponential phase (10). Collectively, these findings substantiate the diverse and complex relations found between bacteria and phagotrophic and photosynthetic protists.

### Bacteria utilizing amoeba-derived nutrients under low-nutrient conditions

SKA58 was the most notable example of a bacterium that could utilize the protists to gain nutrients for proliferation under low-nutrient conditions. Interestingly, in nutrient rich medium, SKA58 was also capable of lysing *A. polyphaga* although at the same time a fraction of the amoeba cells remained viable and dividing while inhibiting bacterial growth. Under low-nutrient conditions, however, SKA58 could grow in co-culture and assimilate amoeba-derived nutrients, probably by lysing amoeba cells (as indicated by the temporary drop in viable amoeba despite bacteria being grazed to low levels). The survival efficacy of SKA58 was, however, not dependent on viable amoeba cells, indicating that SKA58 is more likely to utilize the amoeba as a food resource rather than for a symbiotic or commensal interaction. It is also possible that the amoeba cells do not prey on SKA58 as efficiently under low-nutrient conditions as when nutrients are easily accessible, facilitating bacterial consumption of amoeba-derived nutrients. These results indicate a fine balance in “the battle of eating or being eaten”, and that the result of this interaction is strongly affected by nutrient status. Similarly, *Francisella* spp. display enhanced survival in low-nutrient media when co-cultured with *A. polyphaga* or *A. castellanii* (24, 25), possibly as a result of amoeba-excreted compounds (25). If displayed in natural aquatic environments, the combination of using a protist species to gain nutrients when no other nutrients are available and the ability to lyse the same protist, would indicate a largely unexplored flow of nutrients directly from eukaryotes to bacteria.

### Bacteria requiring viable amoeba to proliferate

We were intrigued to find that six of our studied bacteria species were dependent on viable amoebae to grow but were unable to proliferate in nutrient-rich medium in monoculture (e.g., SKA34). These bacteria possibly proliferate intracellularly, as they were observed within *A. polyphaga*. Notably, none of these strains caused amoeba lysis at 1:1000 bacteria:amoeba ratio, which was used for this experiment, suggesting an endosymbiotic relationship. Four of these bacteria required live amoebas to proliferate which suggest that their growth was more likely stimulated by amoeba-secreted compounds than by consumption of amoeba cell structures or content. Non-lytic, intracellular obligate bacteria within various amoeba species have been documented (2, 6, 45, 49). Bacteria may eventually be packed in vesicles and ejected from their hosts (22, 50), which could explain their presence in the extracellular environment. It is also possible that bacteria can escape grazing and grow extracellularly, supported by compounds secreted by the amoebae that facilitates growth (25). The latter strategy may be akin to ‘microbial gardening’ where protists deliberately enhance bacterial growth to increase their own food supply (51).

### Bacterial-mediated lysis of protists may be linked to type III-VI secretion systems

Bacterial capability to escape predation, lyse protists, and benefit from co-culturing with protists could in many cases be caused by expression of well-known virulence gene homologues, which have been shown to be widespread in marine bacteria, including many of the bacteria in our study (29). As an example, type VI secretion system (T6SS) genes were found in BAL199, SKA14, SKA34, MED193, MED222, MED297, AND4 and RED65 – all capable of causing lysis and/or interacting in some way favorably with amoebae. T6SS is strongly associated with virulence, but T6SS loci are also highly conserved in non-pathogenic bacteria, suggesting other functions than pathogenesis; this could include commensal relationships between bacteria and eukaryotes and social interactions between bacteria (52). In a recent study, T6SSs of pathogenic *Fransciella* spp. did not affect the interaction with various *Acanthamoeba* spp. (25). Isolate MED121 of our study is an example of this type of non-pathogenic, non-dependency relationship with protists. Similar to the type VI secretion system, type III-V (T3SS-T5SS) systems are also associated with bacterial virulence and eukaryotic infections (53). While not as prevalent as T6SS, six strains used in this study possess one or more of these systems (29). Except for MED193, all of these strains caused lysis. In contrast, MED134, BAL38, BAL39 were capable of lysing protists without carrying any known virulence gene homologs (29). However, it should be emphasized that there may also be other unknown virulence genes present in the genomes of these bacteria, which were not included in the Persson et al. (2009) study. Nevertheless, we find it likely that widespread virulence gene homologs in marine bacteria in many cases may have a primary function in commensal or virulent interactions with bacterivorous protists. The pathogenic effects in humans associated with these genes have likely evolved long prior to human contact, perhaps as a result of interactions with protists, as indicated by the presence of T3SS in the predicted chlamydiae ancestor about 700 million years ago (54). The finding that the primary function of T3SSs in the environmental and opportunistic pathogen *Pseudomonas aeruginosa* is to kill biofilm-associated amoebae supports this hypothesis (55). Among the secretion systems, most research has focused on T3SS, likely because of its often found association with disease outbreaks of aquatic animals (31). Incidentally, both bacteria that harbored T3SS were able to induce lysis in two (AND4) or all three (SKA58) protists.

It’s been suggested that protists act as “training grounds’’ for pathogenic bacteria, which then go on to infect secondary species such as humans (56). Alveolar macrophages, in particular, bear many similarities to phagotrophic protists and are susceptible to infection by pathogenic bacteria, which possibly contributes to the symptoms in respiratory diseases (15). This phenomenon has also been described as “coincidental evolution”, and it’s been shown that grazing-resistant *E. coli* are more likely to harbor virulence factors (57). Thus, because neither horizontal nor vertical transmission occurs for many pathogenic bacteria in humans, coincidental evolution may explain their prevalence in secondary hosts. Commonly cited examples of pathogenic bacteria associated with, specifically, *Acanthamoeba* spp., include *Legionella* spp. and *Mycobacteria* spp. (14). These pathogens display enhanced infectivity when affiliated with amoebas (14). Therefore, a better understanding of protist-bacteria interactions may be of pathological and epidemiological importance to other secondary species than just the primary protist.

### Conclusions

In conclusion, our results indicate that marine bacteria may be more commonly capable of escaping protist grazing and, in several cases, even kill protists, than previously recognized. In addition, some of these bacteria may also benefit from, or even require, interactions with viable protists in order to thrive and survive. To mediate bacteria-favorable interactions between bacteria and unicellular eukaryotes may be one of the primary functions of widespread virulence genes in marine bacteria. These complex interactions are likely to be influenced by environmental factors (e.g., temperature and nutritional status) and the physiological status and abundance of the bacteria or protists. Given the ubiquity of temporal dynamics in the composition of natural bacterial and protist communities, the diversity of interactions uncovered here has important implications for how the strength and direction of interactions between bacteria and protists evolve. For example, one could envision a scenario where a sudden release of organic material would trigger the growth of a bacterial population with high capacity to lyse the dominant protists at the time, which would potentially alter the overall grazing pressure on the bacterial community. Thus, the ecological consequences, evolutionary effects, as well as potentially pathogenic and epidemiological potentials of these interactions need to be further elucidated.

## MATERIALS AND METHODS

### Bacteria strains

Bacterial strains of marine origin from the Linnaeus University culture collection, Kalmar, Sweden, were cultured either in liquid medium or on agar plates, dependent on experiment, constituted of three different media. Marine broth 2216 (MB) (Difco) was used for culturing of MED92, MED121, MED134, MED152, MED193, MED217, MED222, MED297, AND4, RED65 and SCB49, ZoBell with Skagerrak seawater (SKA) for SKA14, SKA34, SKA53 and SKA58, and ZoBell with Baltic seawater (BAL) for BAL38, BAL39 and BAL199. Marine agar 2216 (Difco) was used for strains grown in Marine broth. SKA and BAL media were prepared with ZoBell broth, 5 g of peptone (Bacto Peptone; BD) and 1 g of yeast extract (Bacto Yeast Extract; Difco) in 800 ml of Skagerrak or 800 ml Baltic seawater, and 200 ml MilliQ water. For agar plates, 15 g of agar (Difco Bacto Agar) were added to 1000 ml of medium (SKA and BAL). For estimation of growth rates, one colony of each of the 18 bacteria species was harvested from agar plates and added to 50 ml of MB, SKA or BAL. Samples were incubated at room temperature (RT) and monitored once every day, by measuring optical density (OD). Established growth rates were then used to harvest bacteria in late exponential growth phase.

### Eukaryotic organisms

Three species of unicellular eukaryotic organisms were used in the study: *Acanthamoeba polyphaga* (Linc Ap-1), *Euglena gracilis* (CCAP 1224/5Z) and *Tetrahymena pyriformis* (provided by Agneta Andersson, Umeå University, Sweden). Trophozoites of *A. polyphaga*, were maintained aerobically at 27°C in peptone yeast glucose (PYG) medium, at the bottom of 25 cm^2^ culture flasks, as previously described (Axelsson-Olsson, Ellstrom et al. 2007)*. T. pyriformis* was grown in peptone-yeast medium (PPY) at RT in the dark. *E. gracilis* were grown in *Euglena gracilis* Medium 1:1 with Jaworski’s Medium (EG:JM) in a north facing window, according to the manuals from CCAP. For the different experiments, eukaryotic species were grown to a density of 10^6^ cells/ml and then inoculated either in 12-well culture plates or added to 5 ml glass tubes for experiments measuring optical density.

### Bacterial co-culturing with unicellular eukaryotes

Protista species were grown in respective protist media and inoculated with the 18 different bacteria species grown in respective liquid bacterial media at a high (10:1 bacteria:protista) and low (1:1000 bacteria:protista) bacterial ratio using 12-well culture plates to observe consequences of co-cultivation on protists. Additional 12-well plates were inoculated with high bacteria:protista ratio (10:1) to estimate the survival of bacteria in protist co-culture. Cultures of only the different unicellular eukaryotic organisms or bacteria alone in protist media maintained in separate plates were used as controls. Co-cultures and controls were incubated at RT for 15 days. Daily observations of co-cultures with invert microscopy were used to determine protista cell morphology and viability. Moreover, samples were taken every third day and plated on agar plates for the presence of live and growing bacterial cells, to estimate bacterial survival expressed as growth or no growth. All cultures were performed in triplicates. Results were presented as average survival time based on two independent experiments. Observations of intracellular bacterial location in *A. polyphaga* was performed using phase contrast microscopy, and differentiation was made between bacterial presence in vacuoles or in cytoplasm.

### Bacterial survival with A. polyphaga

Bacteria were subjected to three different treatments in PYG media: co-culturing with live amoebae, heat-killed amoebae, and a heat-killed suspension of an earlier co-culture with the same bacteria species. As control, bacteria were grown in PYG medium alone. Heat killed *A. polyphaga* pre-culture was harvested at a concentration of 10^6^ cells/ml and 1 ml per sample were put in 1.5 ml tubes and boiled for 10 min. Heat killed pre-co-cultures were prepared by adding 100 μl bacteria solution with a concentration of 10^4^ cells/ml to 12-well amoebae culture plates and left to grow for 48 hours. From these plates, samples of 1 ml were added to 1.5 ml eppendorf tubes and heat killed by boiling samples for 10 min. Live bacteria were harvested from agar plates, diluted in PYG medium to a final concentration of 10^4^ cells/ml and 100 μl were added to 1 ml of amoeba cultures with different pretreatments (10^6^ protist cells) for each sample in 12-well plates. Duplicate samples were incubated at RT and 100 μl fractions were taken after 24 hours and then once every second day for plating on agar plates for bacterial survival test (growth or no growth). Plates were also observed microscopically to see changes in amoebae morphology.

### Bacterial growth during co-culturing with A. polyphaga

Bacterial samples were taken in approximately late exponential growth phase and added to glass tubes with 5 ml PYG or phosphate buffered saline (PBS), and with or without *A. polyphaga*, at a low concentration, 10^3^ cells/ml at a ratio of 1:1000; bacteria:protista. Samples were incubated at RT for 13 days and monitored daily by measuring OD and observing in microscope. Background (media alone or amoeba monocultures, respectively) was subtracted from experimental values. Bacterial growth and survival were confirmed both by microscopic observation and by plating on agar every second day. Amoebae viability in PYG was determined using the trypan blue exclusion test, initially after one day and thereafter every second day. The cells were suspended by brief vortexing and mixed with 0.04% trypan blue (Sigma-Aldrich, St. Louis, MO). The viability was determined by calculating the ratio of stained versus unstained cells in a Bürcher chamber.

## ACKNOWLEDGMENTS

We thank Sabina Arnautovic for technical support during the experiments and Åke Hagström, Lasse Riemann, and José M. González for inspiring discussions on the potential ecological implications of virulence genes in marine bacteria. We thankfully acknowledge Agneta Andersson for providing cultures of *Tetrahymena pyriformis* and *Euglena gracilis* and advice on cultivation conditions. The research was supported by The Swedish Research Council VR and the Swedish strong research environment EcoChange (Ecosystem dynamics in the Baltic Sea in a changing climate).

## SUPPORTING INFORMATION

Additional Supporting Information may be found in the online version of this article at the publisher’s website.

---

**Table S1.** Bacterial survival of all strains included in the study measured for a maximum of 19 days in monoculture and co-culture with amoeba in different settings. Bacteria were cultured in the presence of viable *A. polyphaga* (co-culture), heat-inactivated *A. polyphaga* or a heat-inactivated previous co-culture of *A. polyphaga* and the same bacterial strain and compared to bacteria growing in monoculture. The survival time in days of independent duplicate samples are shown as X/Y.

| Bacterial strain | Co-culture<br>(days) | Heat inactivated<br>amoeba (days) | Heat inactivated co-culture (days) | Bacterial mono-<br>culture (days) |
| --- | --- | --- | --- | --- |
| MED92 | 3/3 | 3/3 | 3/3 | 3/3 |
| MED121 | 3/3 | 3/3 | 3/3 | 5/3 |
| MED134 | 13/15 | 5/5 | 5/5 | 5/7 |
| MED152 | 1/3 | 3/3 | 1/3 | 3/3 |
| MED193 | 13/15 | 5/5 | 5/5 | 5/5 |
| MED217 | 17/15 | 15/15 | 15/15 | 17/15 |
| MED222 | 19/19 | 5/5 | 7/5 | 7/11 |
| MED297 | 15/15 | 11/15 | 15/15 | 15/15 |
| AND4 | 19/19 | 11/15 | 11/9 | 11/9 |
| RED65 | 9/7 | 5/5 | 9/9 | 5/5 |
| SCB49 | 17/15 | 15/15 | 15/15 | 15/15 |
| BAL38 | 19/13 | 19/19 | 19/19 | 19/19 |
| BAL39 | 19/19 | 19/19 | 19/19 | 19/19 |
| BAL199 | 15/15 | 13/15 | 13/15 | 3/15 |
| SKA14 | 19/19 | 19/19 | 19/19 | 19/19 |
| SKA34 | 19/19 | 5/5 | 3/0 | 7/5 |
| SKA53 | 13/15 | 13/15 | 13/15 | 15/15 |
| SKA58 | 19/19 | 19/19 | 19/19 | 19/19 |

**Fig S1.**
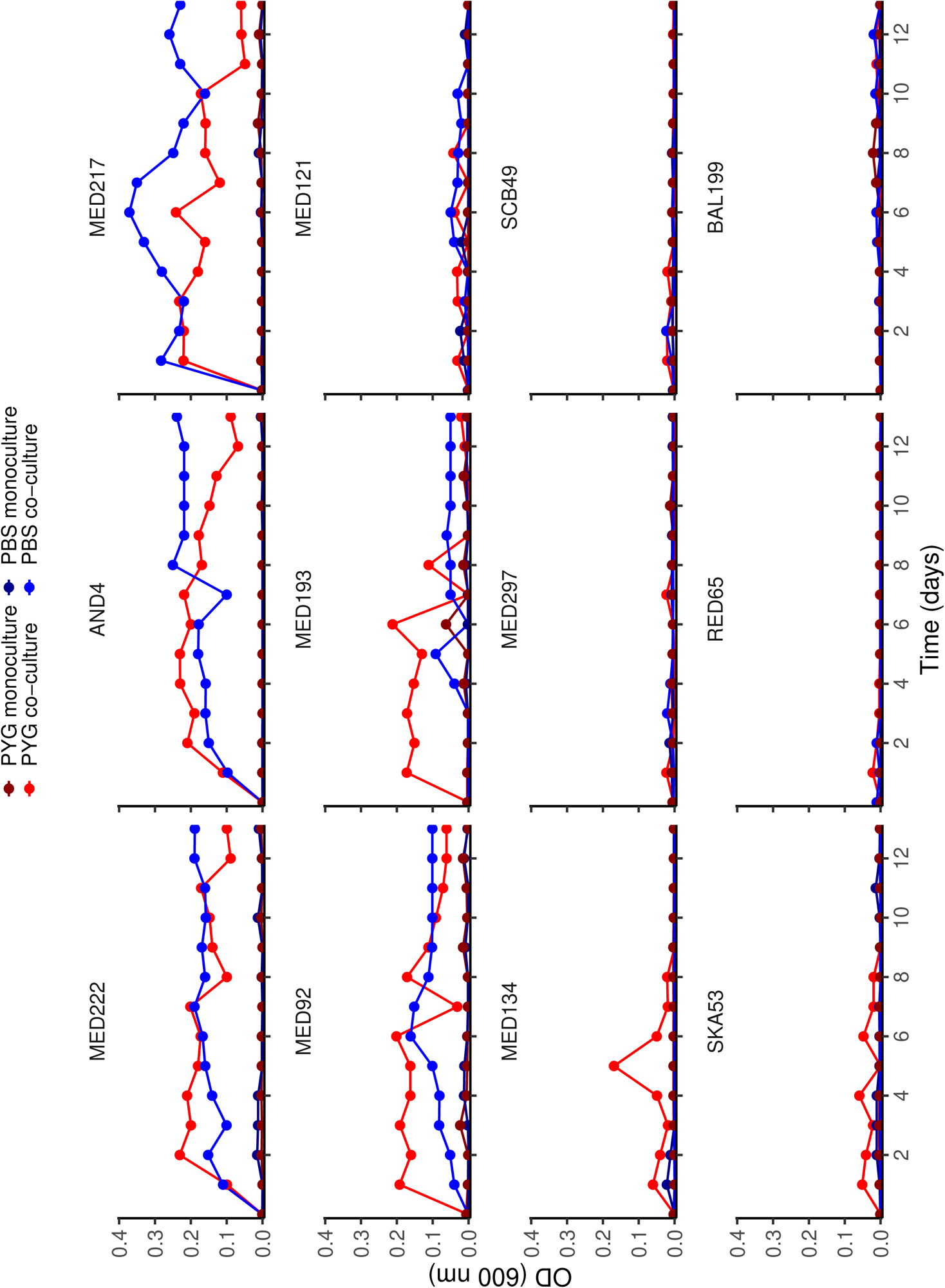
Growth curves of the remaining 12 marine bacterial strains not included in Fig. 3. The bacteria were cultivated in nutrient-rich media (peptone yeast glucose [PYG]), low-nutrient media (phosphate buffered saline [PBS]), and in mono- and co-culture with *A. polyphaga*. All bacteria cultures started from late exponential growth phase and the growth was determined by plating.

